# Functional Connectivity Alterations in Major Depressive Disorder

**DOI:** 10.64898/2026.03.07.710303

**Authors:** Malvika. Sridhar, Sir-Lord Wiafe, Bradley Baker, Vince. D. Calhoun

## Abstract

Major depressive disorder (MDD) is associated with brain-wide network disruptions. This study investigates a large resting-state functional magnetic resonance imaging dataset (N = 519) to analyze static and dynamic functional network connectivity (FNC). Using independent component analysis, our analysis revealed hyperconnectivity within sensorimotor and temporal subdomains, hypoconnectivity from higher cognitive networks, and hyperconnectivity from the default mode and sensorimotor domains in MDD. A novel frequency-sensitive dynamic approach identified disruptions in the temporal synchrony of brain states engaging the default mode-paralimbic, sensorimotor, and frontal regions, as well as the subcortical limbic, frontal, and salience regions. Overall, the findings highlight the utility of combining static and dynamic approaches in large neuroimaging datasets to elucidate the neural underpinnings of MDD pathology.

## 1. INTRODUCTION

Major depressive disorder (MDD) is a highly debilitating psychiatric disease characterized by changes in mood, behavior, and cognition. Resting-state functional magnetic resonance imaging (rs-fMRI) studies consistently indicate disrupted functional connectivity (FC) within and between large-scale brain networks, particularly the default mode (DMN), central executive (CEN), and dorsal attention (DAN) networks [1], [2]. Hyperconnectivity within the DMN and the DMN-DAN is thought to underlie maladaptive self-referential processing and rumination bias, while reduced CEN and CEN-DAN connectivity is linked to poor cognitive-affective processing [1].

FC typically assesses coherent activity of low-frequency blood-oxygen-level-dependent (BOLD) signal fluctuations across spatially distinct brain regions [4]. Independent component analysis (ICA)-based methods provide a data-driven framework to examine these alterations at the network level, overcoming limitations of region-based approaches [5]. ICA allows the identification of spatially robust and functionally meaningful intrinsic connectivity networks (ICNs) without prior anatomical constraints. Importantly, the ICNs can overlap with one another, separating artifacts from the signal and distinguishing components of multi-functional regions.

While FC has revealed useful insights into brain function, dynamic FC offers a richer, temporal view of neural coupling by capturing moment-to-moment spatiotemporal dynamics of brain networks [6] and has identified reduced network flexibility and altered temporal stability in MDD [6], [7]. Dynamic functional network connectivity further extends this framework to characterize interactions between ICNs identified by ICA, with studies reporting prolonged dwell times in weakly connected states in MDD [8].

Predominantly researched on Western populations, findings are limited by cross-cultural validity [9]. Recent studies in China found reduced DMN connectivity in MDD linked to depressive severity, sensorimotor network (SMN) alterations [10], [11], and biological subtypes with opposing patterns of network connectivity [12]. With a need for population-specific analyses, the primary aim of this study is to investigate static and dynamic functional network connectivity (sFNC and dFNC) abnormalities in MDD within a large rs-fMRI Chinese dataset using spatially constrained ICA (sc-ICA). Importantly, we analyzed dynamic connectivity using the recently developed single-sideband modulation sliding-window Pearson correlation (SSB-SWPC) [13]. This preserves the full spectral information of the BOLD signal (0.01-0.15 Hz) and enables higher temporal precision and sensitivity to transient connectivity patterns compared to traditional dFNC methods.

## 2. METHODS

### 2.1. Participant*s*

519 Han Chinese participants were recruited from four hospitals in China (Table 1). All patients were confirmed by the DSM-IV. The current depression severity was rated using the 17-item Hamilton Depression Rating Scale (HDRS). Ethical approval and informed consent were obtained, per the Institutional Review Boards.

**Table 1.** Demographics.

|  | Sample | Age | Sex (M: F) | HDRS |
| --- | --- | --- | --- | --- |
| MDD | 235 | 32.289 ± 10.899 | 93:142 | 20.497 ± 6.146 |
| HC | 284 | 31.320 ± 10.696 | 104:180 | N/A |

**Abbreviations:** MDD, Major Depressive Disorder; HC, healthy controls; M, Male; F, Female; HDRS, Hamilton Depression Rating Scale

### 2.2. MRI Acquisition

All participants were screened for MRI safety before scanning procedures. At all four sites, a single 8-minute resting state scan was obtained using a 3T scanner. A total of 240 volumes of echo-planar images were obtained (repetition time/echo time (TR/TE) = 2,000/30 ms, field of view (FOV) = 64 × 64 matrix, flip angle (FA) = 90°). All participants were instructed to keep their heads still and stay awake with their eyes closed during the scan. Memory foam and inflatable padding were used to restrict head motion.

### 2.3 Preprocessing

All scans were preprocessed using the SPM12 (fil.ion.ucl.ac.uk/spm/) software. An automated pipeline developed at the Brainnetome Center (brainnetome.org) was used. This included the removal of the first 10 volumes to account for T1 equilibration effects, slice timing and motion correction, spatial normalization to the Montreal Neurological Institute template, spatial smoothing (8-mm-full-width, half-maximal Gaussian kernel), and regressing non-brain tissue signals, global signal, and 24 motion parameters. Each voxel time series was z-scored to normalize variance across space. After preprocessing, 105 gray matter-based spatial ICNs and their associated time series were derived using sc-ICA with the Neuromark 2.2 template [14]. The 105 ICNs span 7 domains and 14 subdomains as shown in Figure 1. Each ICN was bandpass filtered to a frequency band of [0.01-0.15] Hz to capture relevant neural signals that were then z-scored.

**Fig. 1.**
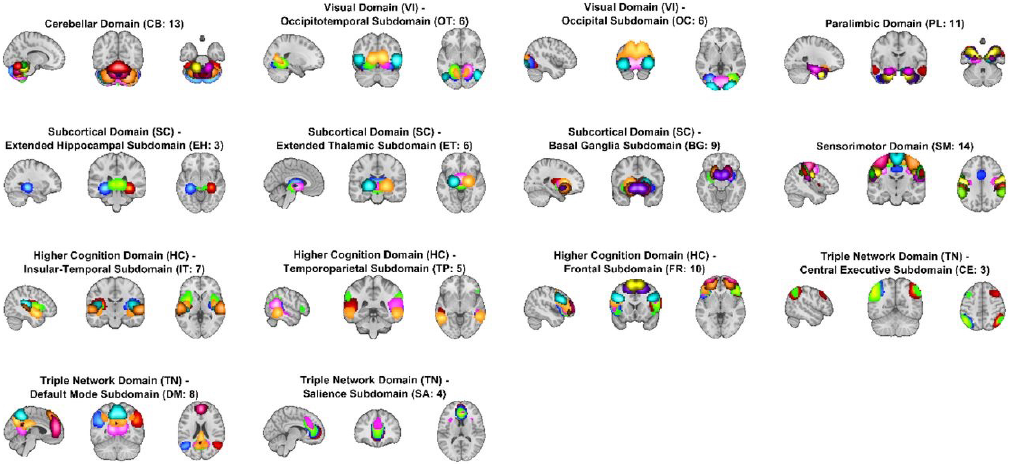
The overlayed spatial maps of the 105 intrinsic connectivity networks from the NeuroMark 2.2 multi-scale template atlas are plotted above (Jensen, Kyle M et al. 2024 [14], under a Creative Commons license CC BY-NC-ND 4.0.

### 2.4 Analyses

To derive sFNC, cosine similarity was computed between time series of ICN pairs. Both groups showed high similarity in FNC across sites (all pairwise Pearson MDD r > 0.91; HC r > 0.88), indicating consistency across acquisition locations. dFNC was computed using SSB-SWPC with a short 44-second (width = 22 TRs) tapered sliding window created by convolving a Gaussian rectangle (σ = 3 TRs) [16]. SSB-SWPC utilizes Hilbert-based frequency modulation to shift the BOLD signal upward in the frequency spectrum, enabling shorter window lengths without compromising low-frequency information during high-pass filtering. Next, blind ICA was used to extract 15 population-level FNC patterns or “states”. Each state reflects overlapping and recurrent FNC patterns that a subject may express simultaneously. The states served as priors in a constrained ICA to estimate 15 individual subject-specific states and their time series (expression of a state at a given time point). Pearson correlation was computed between time series, resulting in 15×15 correlation matrices to capture the degree to which brain states co-fluctuate. Group-level comparisons were then performed on the resulting state co-expression (“synchrony”) patterns. Ordinary least squares regression was used to test group differences, covarying for sex, age, and site. Multiple comparisons were performed using a local false discovery rate (FDR) approach [15], and results with FDR p<0.05 were considered significant.

## 3. RESULTS

### 3.1. sFNC

The regression model resulted in 337 significant ICN pairs between groups (Fig. 2a). The top 53 pairs (FDR p<0.01) have been visualized in a circular connectogram in Fig. 2b. MDD participants show increased FC within higher cognitive temporal and SMN domains, with decreased FC within higher cognitive frontal regions, relative to HCs. MDD participants also showed increased between-domain FC from the SMN to the salience and basal ganglia regions, as well as from the DMN to paralimbic and frontal regions. In contrast, decreased FC is observed from the higher cognitive networks to visual (VI), paralimbic, and salience networks. No significant associations were found between imaging findings and depression severity.

**Fig. 2.**
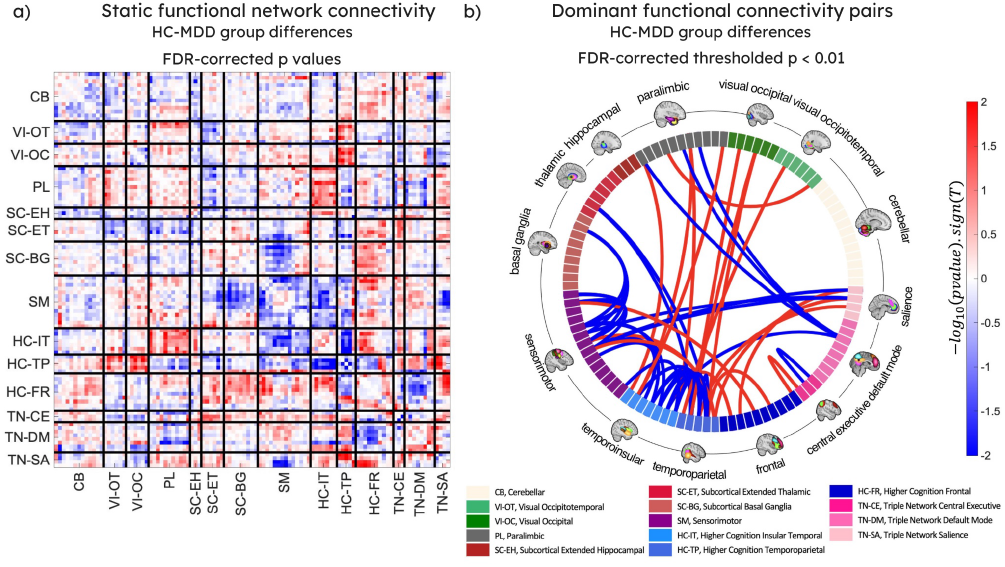
a) FDR-corrected matrix of group differences in sFNC visualized. **b)** Circular plot thresholded to visualize the most dominant sFNC pairs. In terms of connectivity, *blue=MDD*>*HC, red=MDD*<*HC*.

### 3.2. dFNC state synchrony

The results of our group-level comparisons revealed significant differences in the state synchrony of 4 state pairs (Fig. 3a). When compared with healthy controls, participants with MDD show greater synchrony between states 4 and 10, states 14 and 5, and states 14 and 8 (Fig. 3b). Conversely, reduced synchrony was observed between states 4 and 11 (Fig. 3b).

**Fig. 3.**
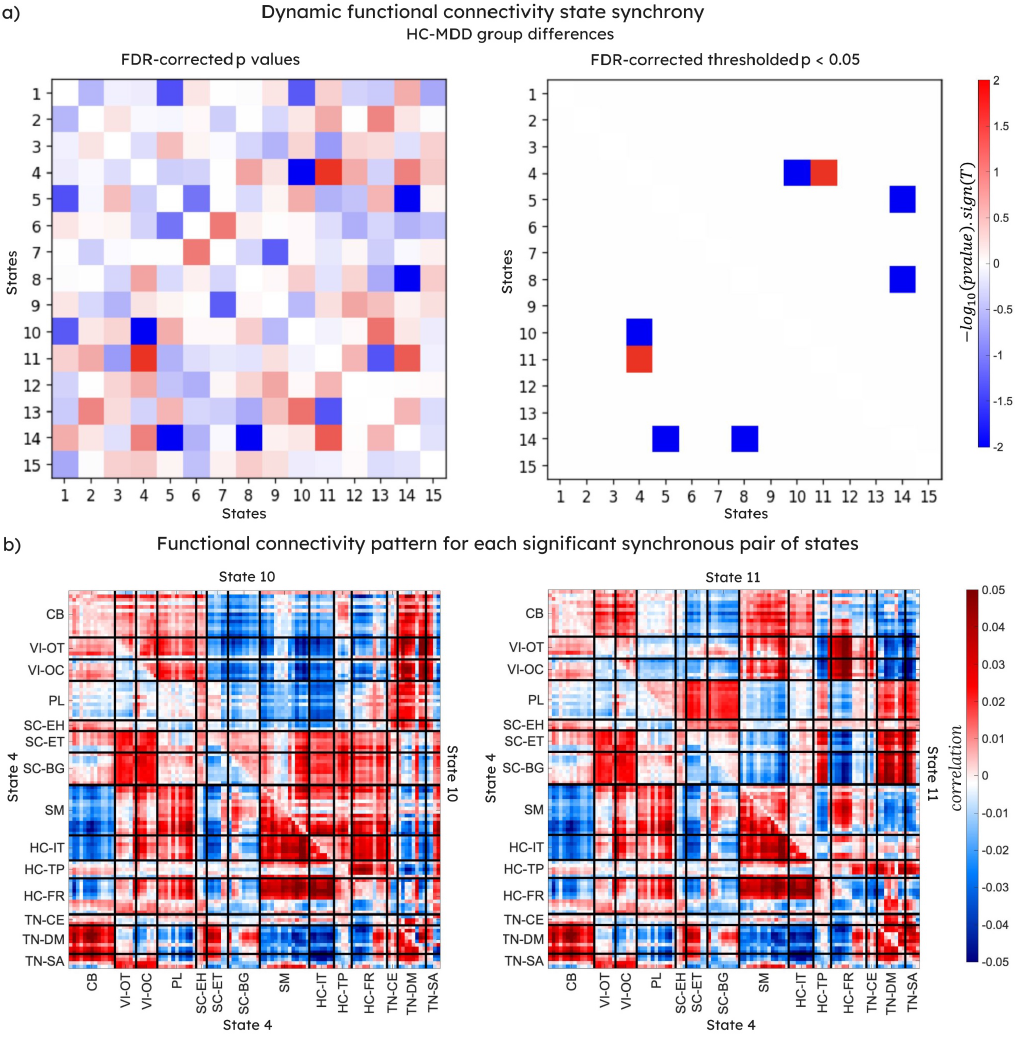
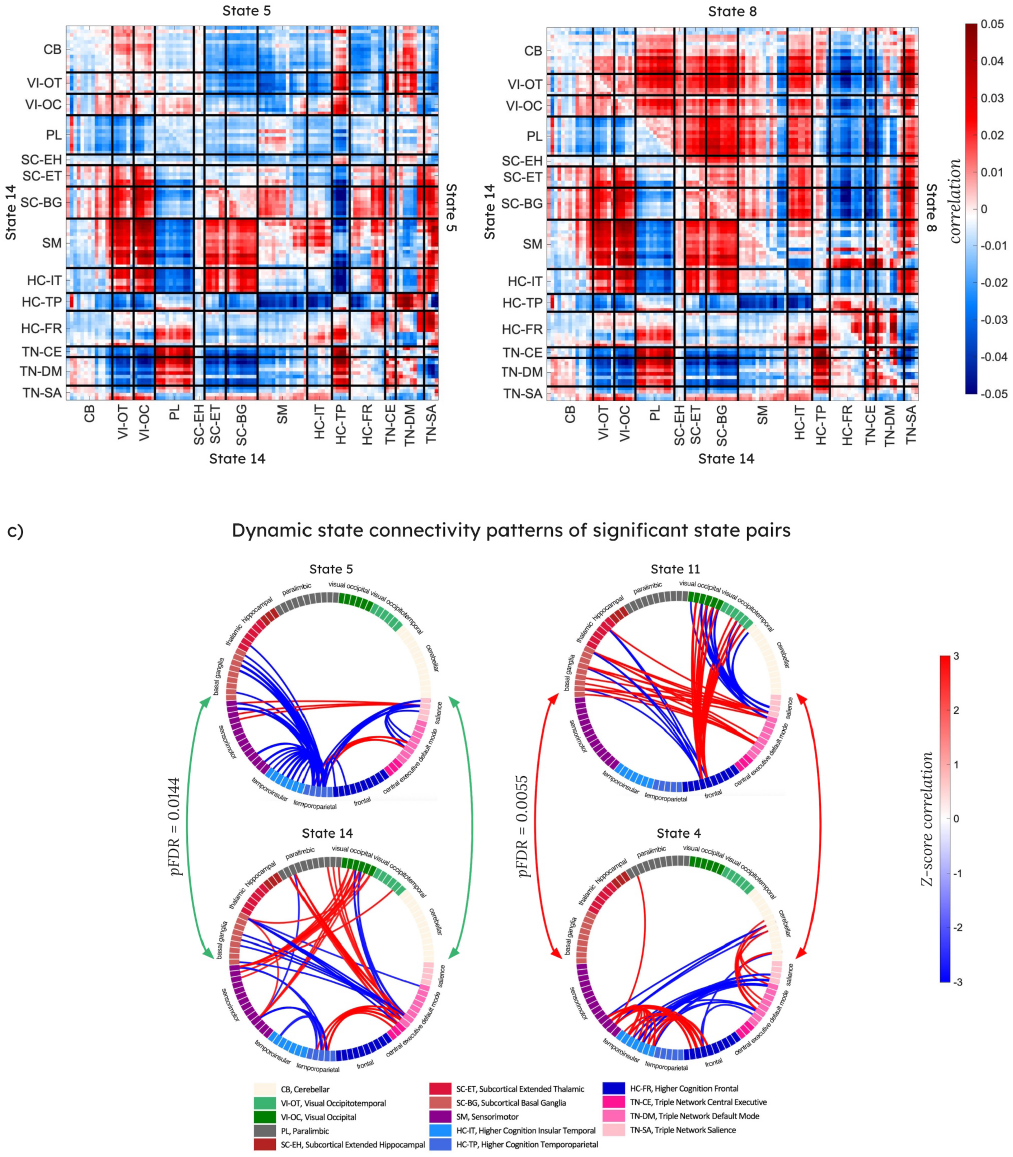
Group differences in the synchrony of dFNC brain states **a)** Image on the left shows FDR-corrected p-values and the right shows values thresholded to pFDR < 0.05. In terms of synchrony, *blue=MDD*>HC, *red=MDD*<*HC* **b)** FNC matrices for all four significant pairs are illustrated. Each matrix represents a state pair with the lower triangular matrix to show one state and the upper, its corresponding pair: state4-state10 (top-left, pFDR = 0.0223*); state4-state11 (top-right, pFDR = 0.0055**); state14-state5 (bottom-left, pFDR = 0.0144*); state14-state8 (bottom-right, pFDR = 0.0195*). **c)** Figure shows member state connectograms for the top 2 significant state synchrony pairs: state14-state5 (in sync) and state4-state11 (out of sync), *blue=reduced FC, red=increased FC*.

The strongest 2 pairs showed increased synchrony in MDD between states 14 and 5 (involving the temporal, basal ganglia, triple network, paralimbic, SM, and VI regions) and decreased synchrony between states 11 and 4 (involving the subcortical limbic (basal ganglia, thalamic) and hippocampal, frontal, visual, salience, insular, DMN and cerebellar regions (Fig. 3c).

## 4. DISCUSSION

Our findings demonstrate widespread differences in FNC patterns within core neurocognitive systems between MDD and healthy participants. sFNC results highlight hyperconnectivity within temporal and SMN regions, and hypoconnectivity within frontal regions in MDD patients. When examining between-domain connectivity, predominantly decreased connectivity is observed from higher cognitive networks. Primarily encompassing the lateral prefrontal cortex, this aligns with findings of general deficits in top-down regulation of attention and emotion and a bias toward rumination at the cost of attending to external stimuli [16], [17].

Contrary to our findings, hypoconnectivity within SMN regions [18] and between SMN and other networks has been observed in previous studies [19]. While this may be attributed to differences in study design or demographics, an intriguing future exploration could delve into the role of the SMN FC as a population and symptom-specific feature of MDD.

We also observed increased FC between DMN and other brain networks, corroborated by prior findings [1], [20], [22]. Still, other studies have noted decreased connectivity from DMN nodes to other brain regions [21]. Overall, Fig. 2a shows decreased between-domain and increased within-domain connectivity in MDD, suggestive of reduced cross-communication and integration across brain networks in the diseased state. The absence of significant associations with HDRS suggests that observed network disruptions may be reflective of chronic trait-like neural markers of MDD rather than current severity. Notably, we do not have access to item-level HDRS scores were unable to examine associations with individual symptom dimensions. Moreover, while participants from site 2 were confirmed to be unmedicated and first-episode MDD, this information was not obtained at the other sites, limiting our ability to control for medication class or illness duration robustly. We acknowledge these as important factors that necessitate future research in similar datasets with detailed clinical information.

Our dFNC results showed significant group differences in the synchrony of specific brain states. MDD participants exhibited increased temporal synchrony between 3 state pairs and decreased synchrony between 1 state pair, suggesting altered coordination in the temporal dynamics of state expression. States 14 and 5 exhibit temporoparietal-DMN hyperconnectivity and temporoparietal-SMN hypoconnectivity. Increased synchrony of these states in MDD may reflect a mode of heightened internal focus, affective salience, driven by concurrent engagement of the DMN, paralimbic, and VI systems, and reduced external sensory integration from the disengagement of salience and temporal higher cognitive systems [16],[23]. The reduced synchrony between state 4, an integral network switching hub of affective-interoceptive processing (frontal-insular and DMN-cerebellar coupling and insular-salience disengagement [27]) and state 11, an emotional stimuli response and regulation system (driven by salience-limbic and frontal-VI coupling and frontal-limbic and salience-VI decoupling [23], [26]) may reflect impaired coordination in MDD. Thus, modeling interstate relationships might be important when characterizing the dynamic architecture of brain function in MDD.

Several limitations should be considered when interpreting these findings. First, detailed information on medication status was unavailable across sites, potentially confounding group-level differences. Yet, a substantial proportion of MDD patients remain treatment-resistant or respond heterogeneously to pharmacological interventions, making it difficult to establish consistent group-level associations between medication class and connectivity patterns. Thus, we believe our analysis still offers meaningful insight into trait-level neural alterations in MDD. Second, we were unable to distinguish between the first episode and recurrent/chronic MDD in the broader sample. Third, we only had access to the total HDRS scores, which limited our ability to examine associations to specific symptom dimensions (e.g., insomnia, appetite, anxiety) using HDRS subscales. Fourth, our results focused on a homogeneous ethnic group and additional analyses is needed to confirm generalizability. Finally, the use of eyes-closed resting-state fMRI may have introduced variability in internal state and arousal levels. Still, prior work showed that drowsiness tends to cluster into specific dynamic states [27], and meaningful fluctuations still occur within non-drowsy states, allowing clinical groups to be distinguished [27]. These findings further underscore the value of dynamic analyses in resting-state fMRI. Future work could replicate these findings in independent samples and explore cross-validation approaches to strengthen the generalizability of ICA-based methods.

## 5. ACKNOWLEDGMENTS

This work was supported in part by NIH R01MH12361 and NSF 2112455.

## Notes

### Competing Interest Statement

The authors have declared no competing interest.

