## Supplemental Figures for "Functional Connectivity Alterations in Major Depressive Disorder"

**Functional Connectivity Alterations in Major Depressive Disorder: A Resting-State fMRI Study**

***Online Supplementary Content***

**Description:** The 15 distinct population-level FNC patterns or “states” were derived using a blind-ICA method performed on the concatenated dFNC matrices of the entire sample. Each state reflects an overlapping and dominant brain connectivity pattern such that a subject may express multiple “states” at a given point in time. These 15 states (visualized in this supplement) were used as priors in a constrained ICA framework to estimate individual subject-level state representations. This data-driven approach enables the analysis of the brain's complex functional nature without oversimplification.

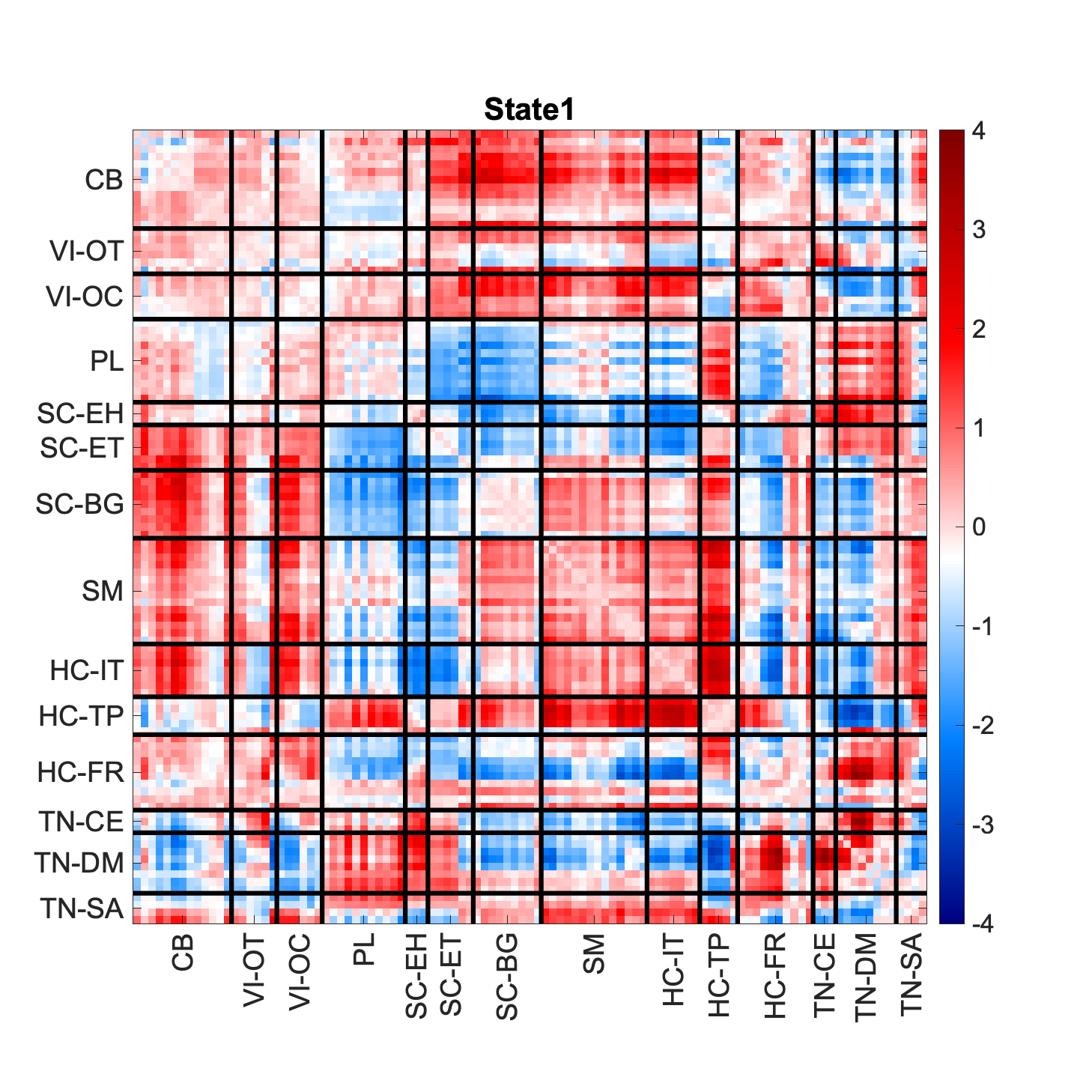

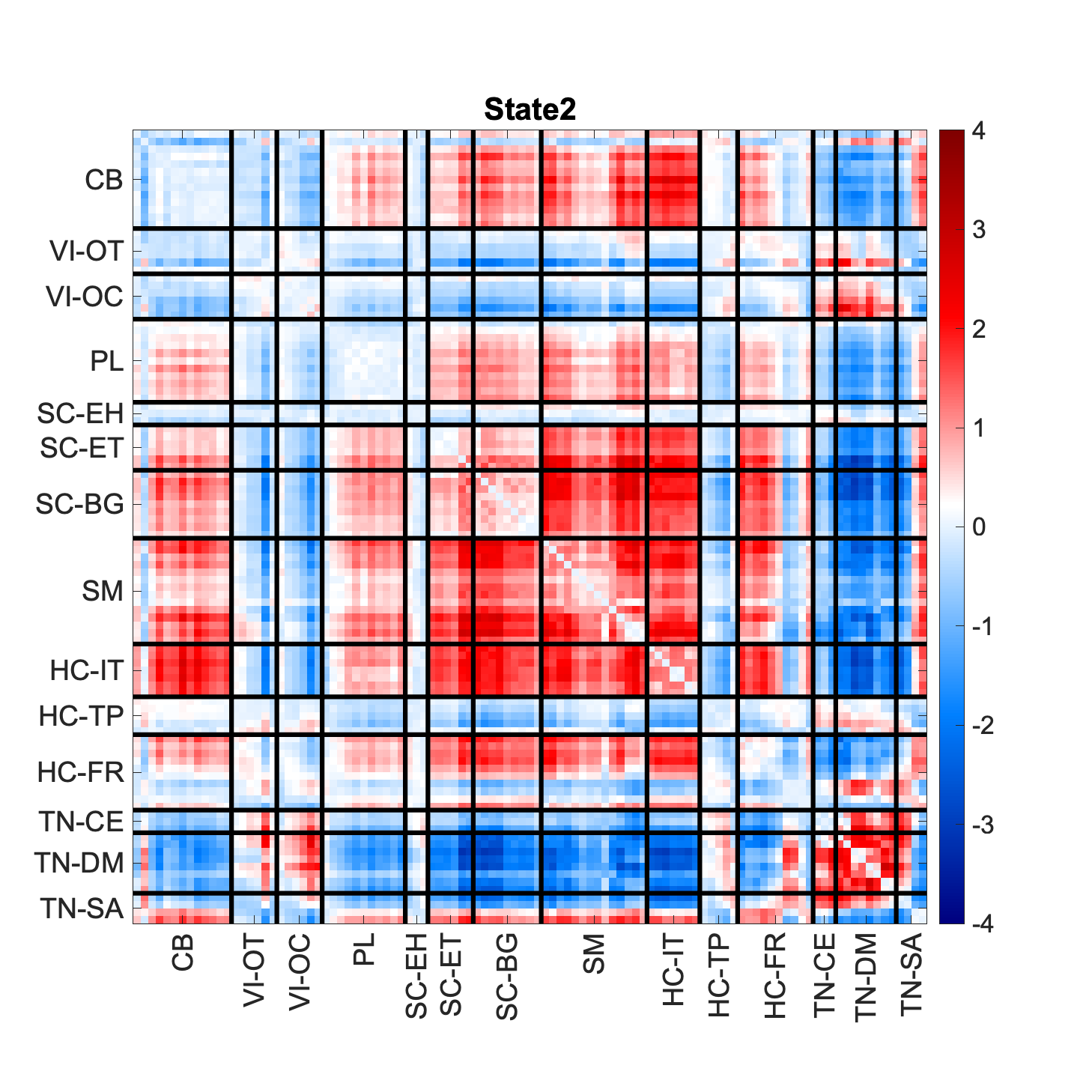

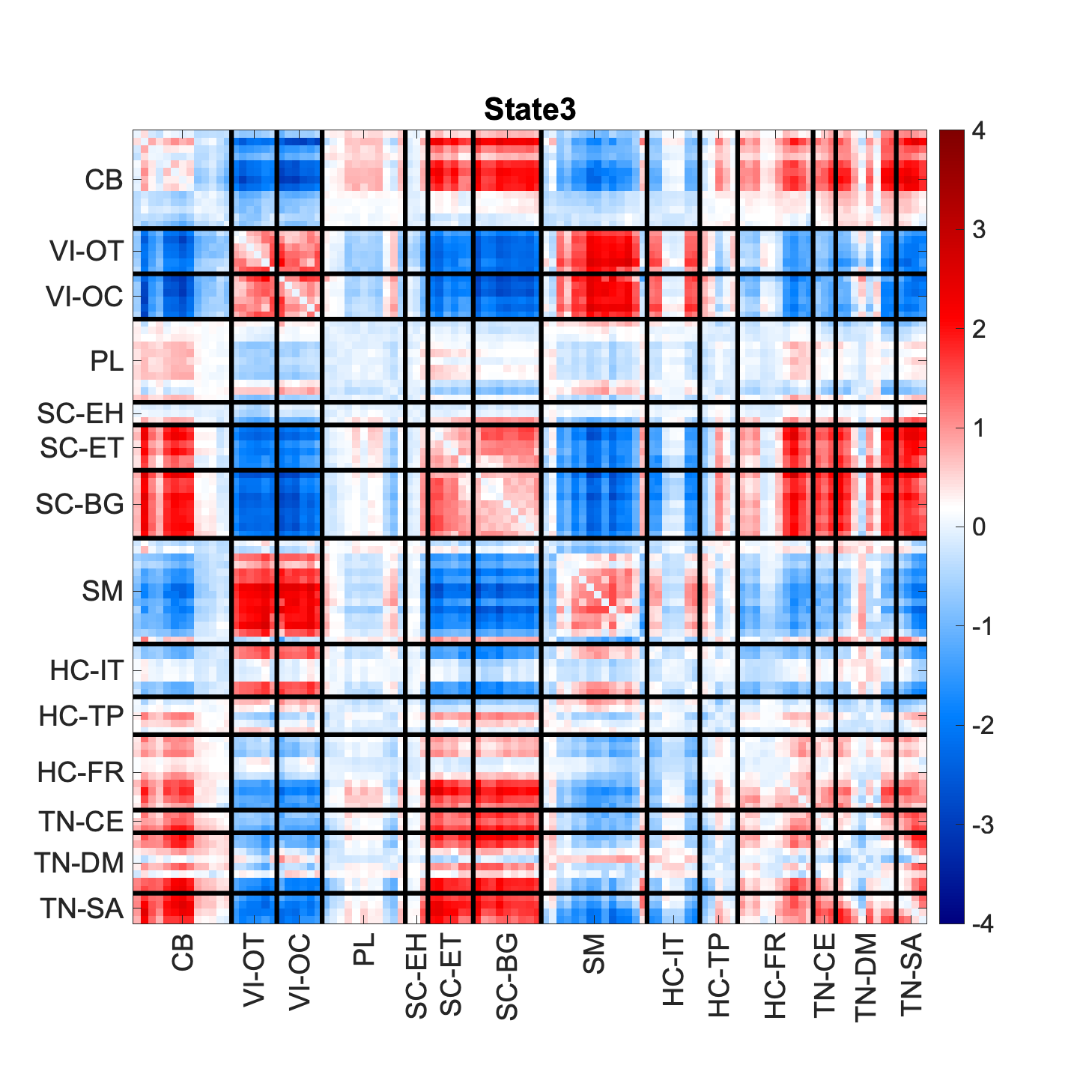

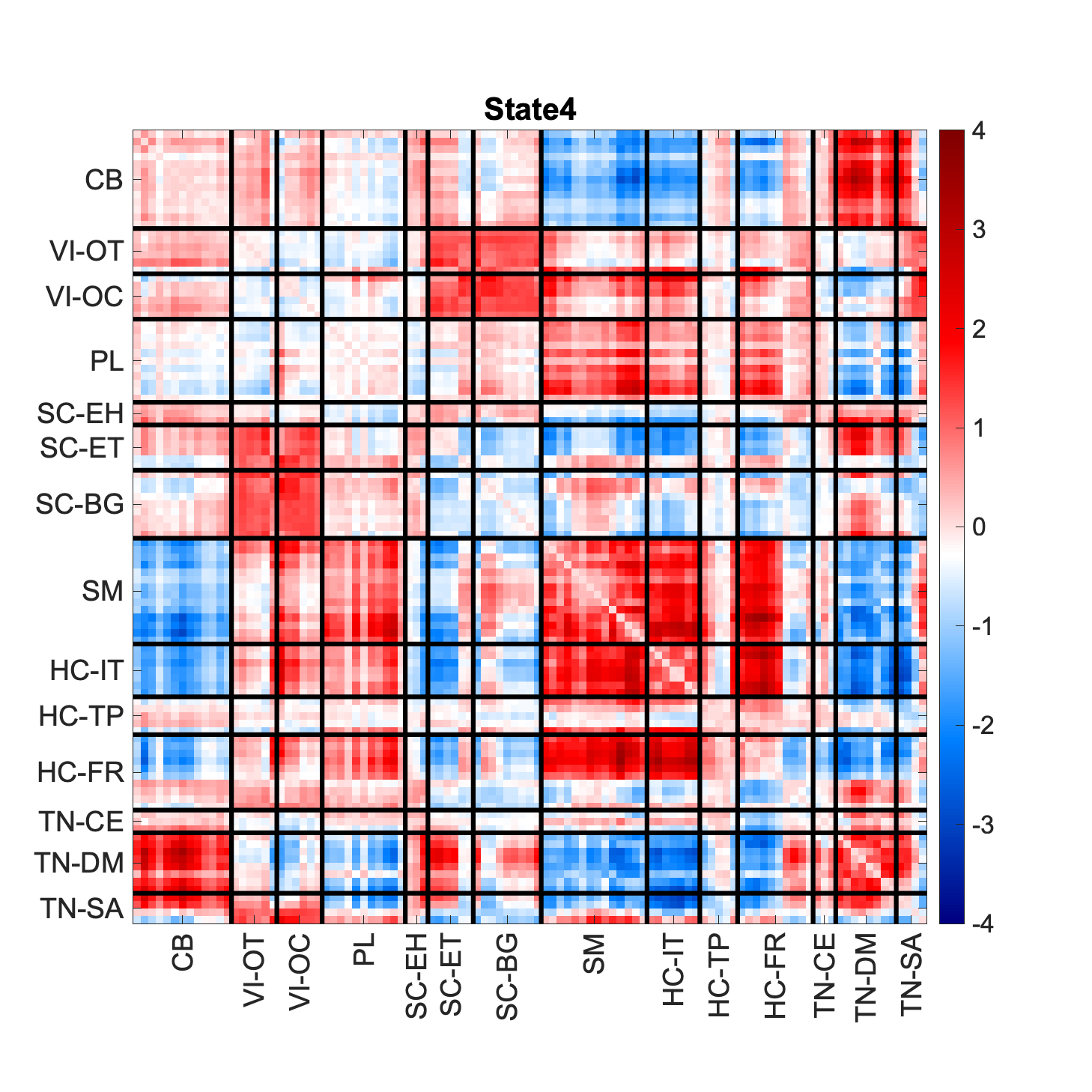

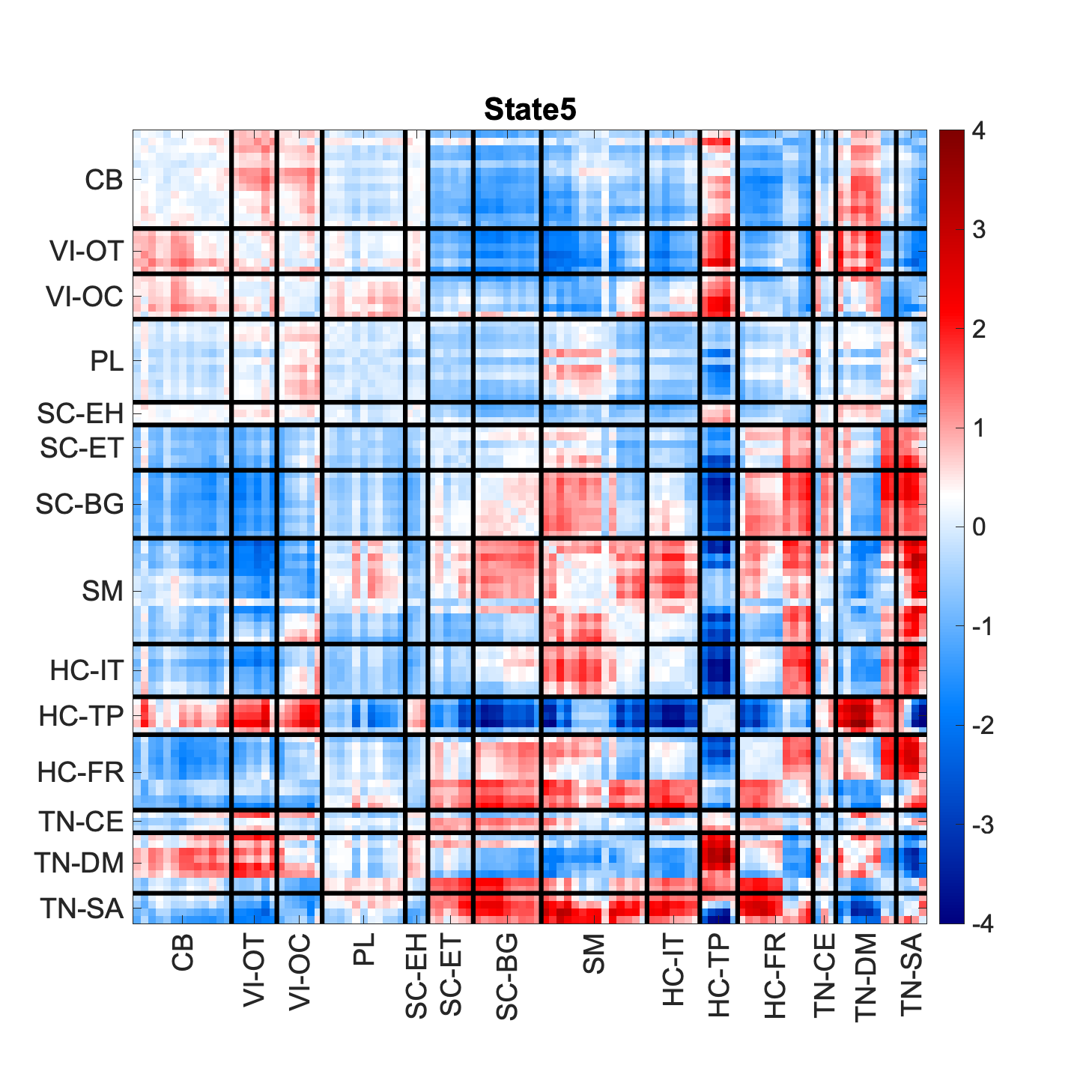

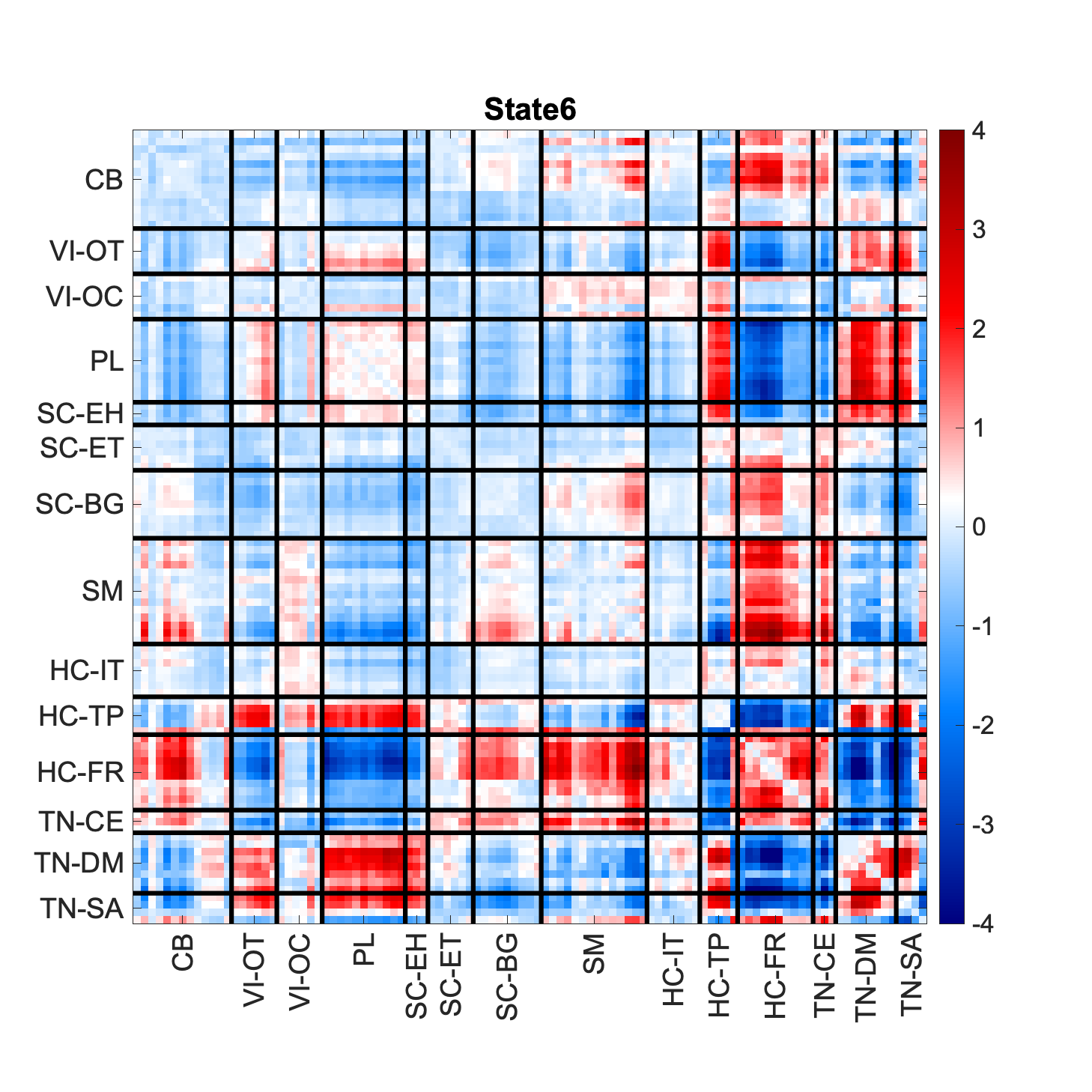

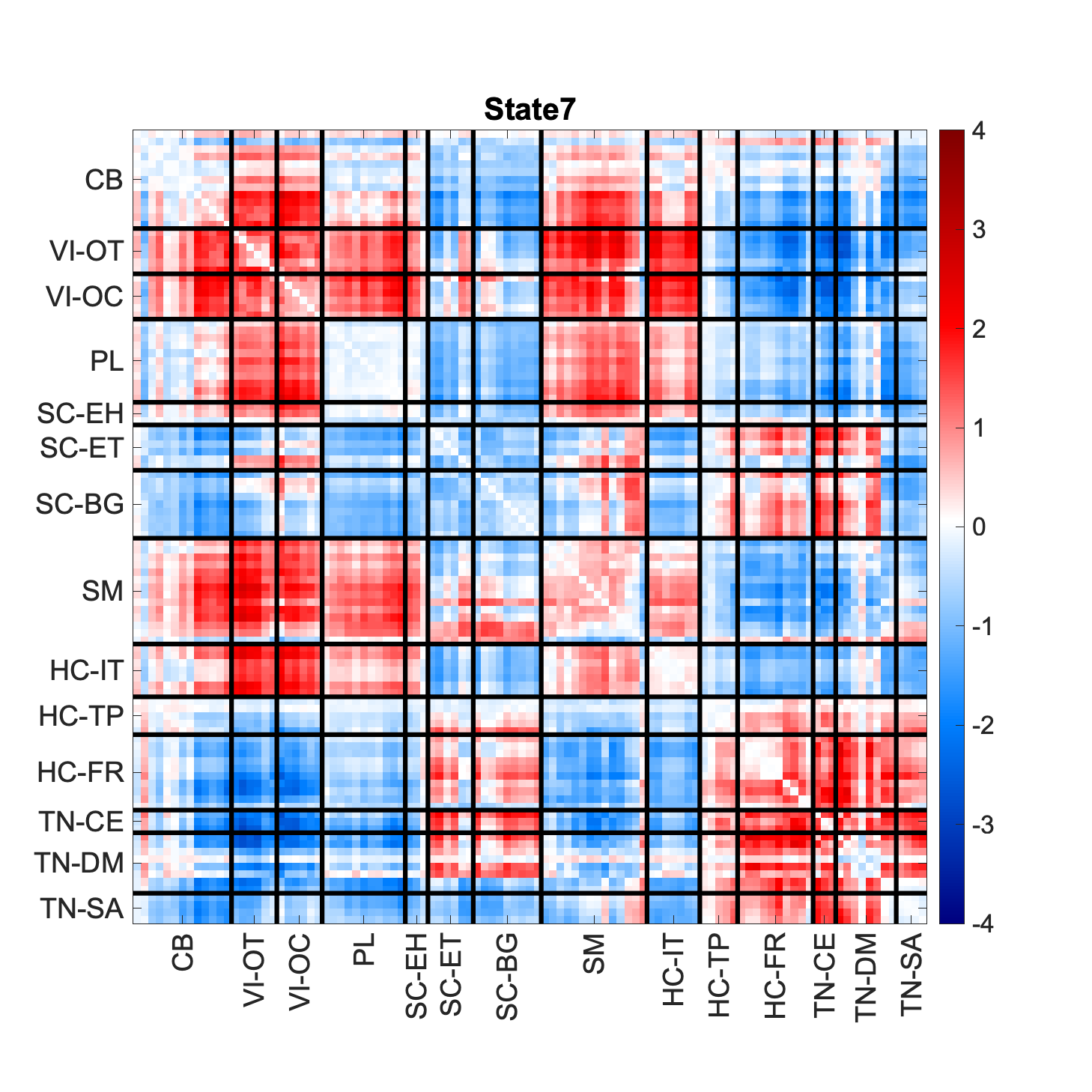

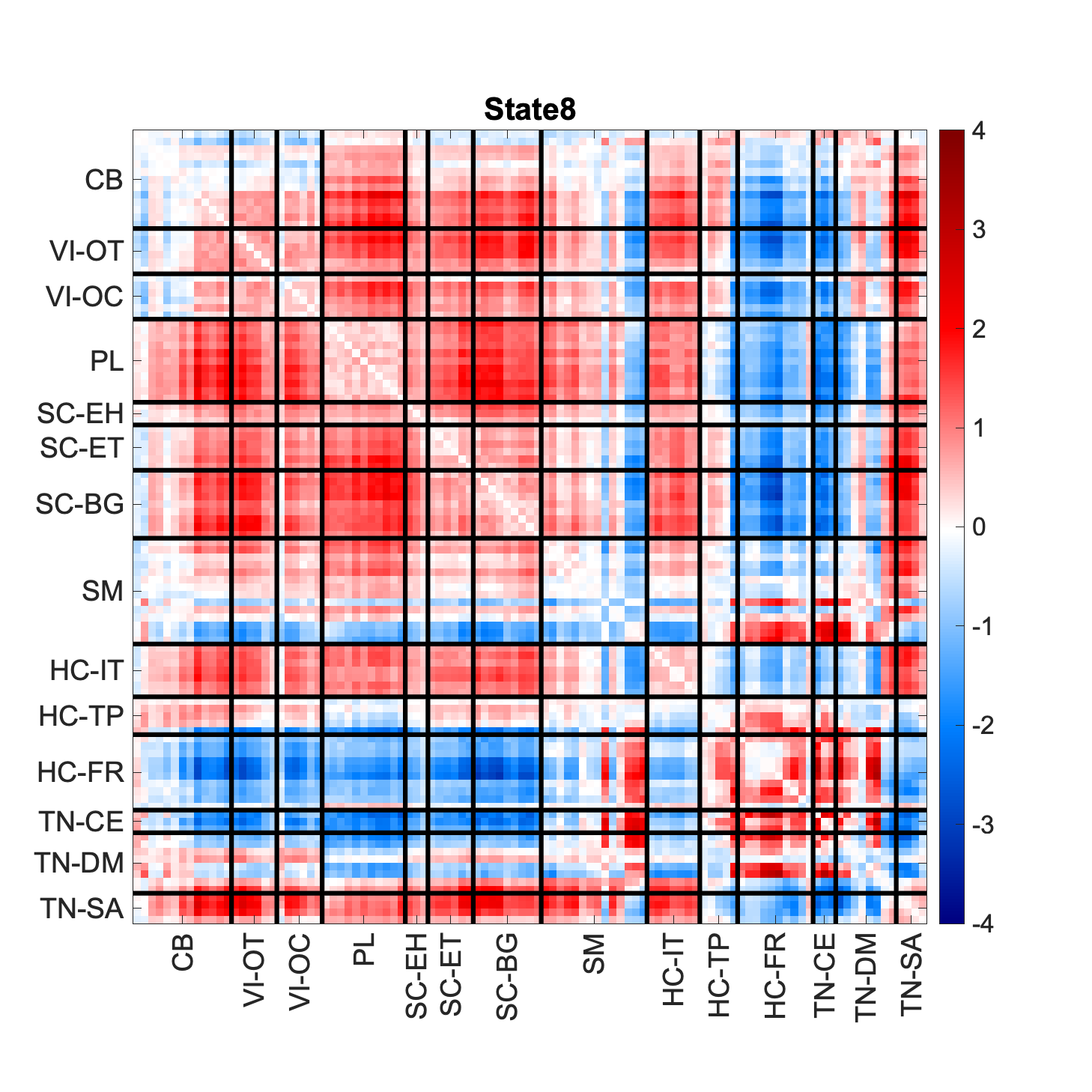

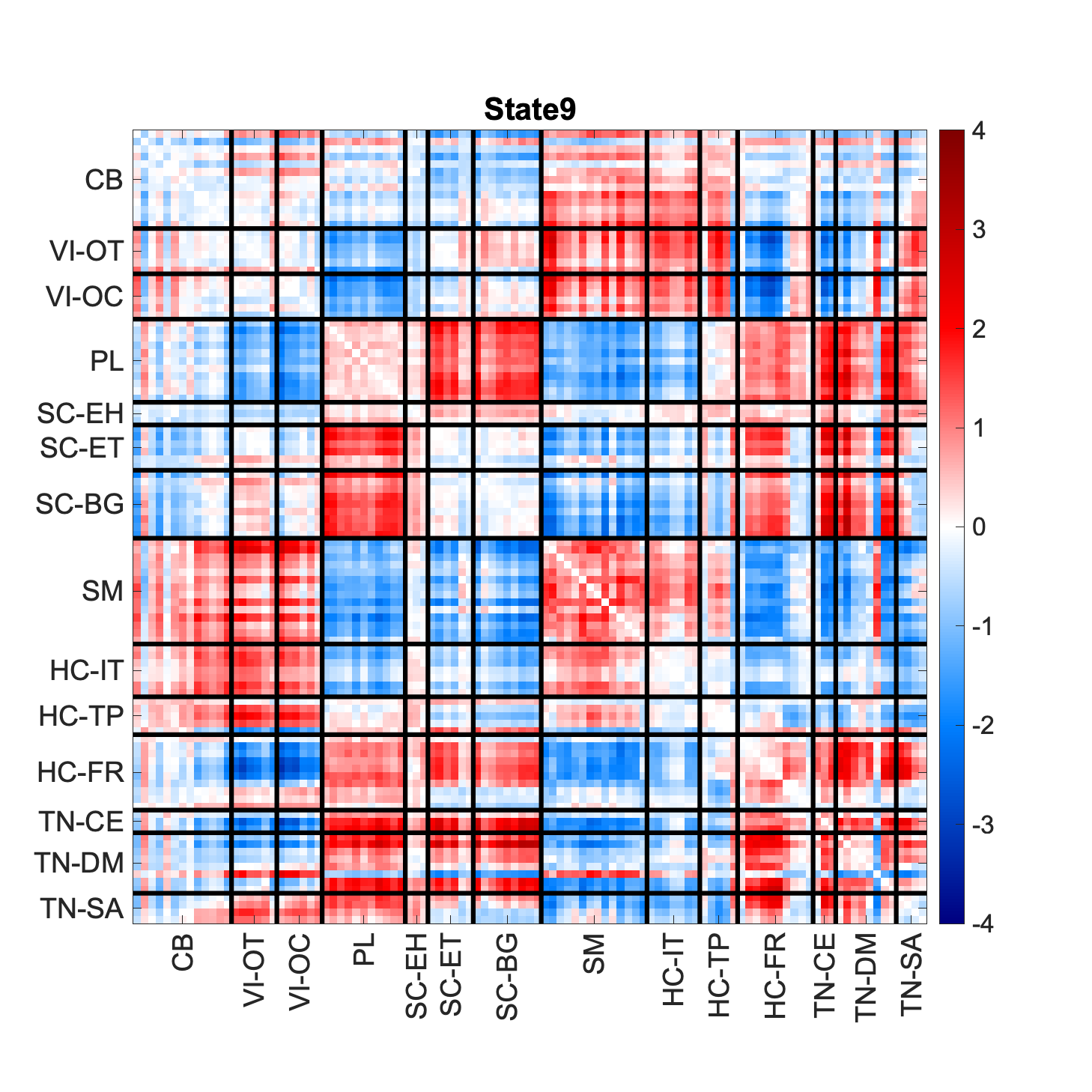

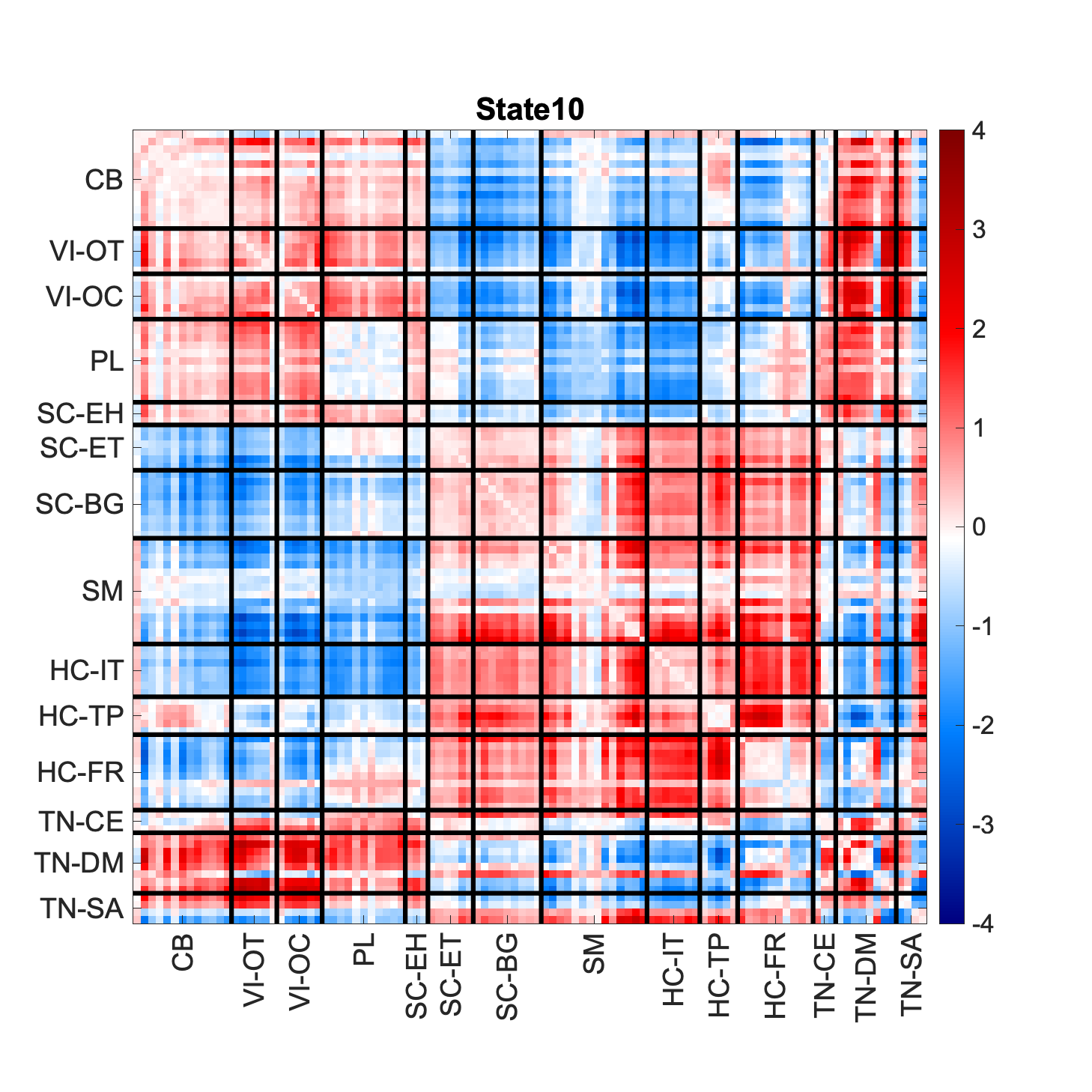

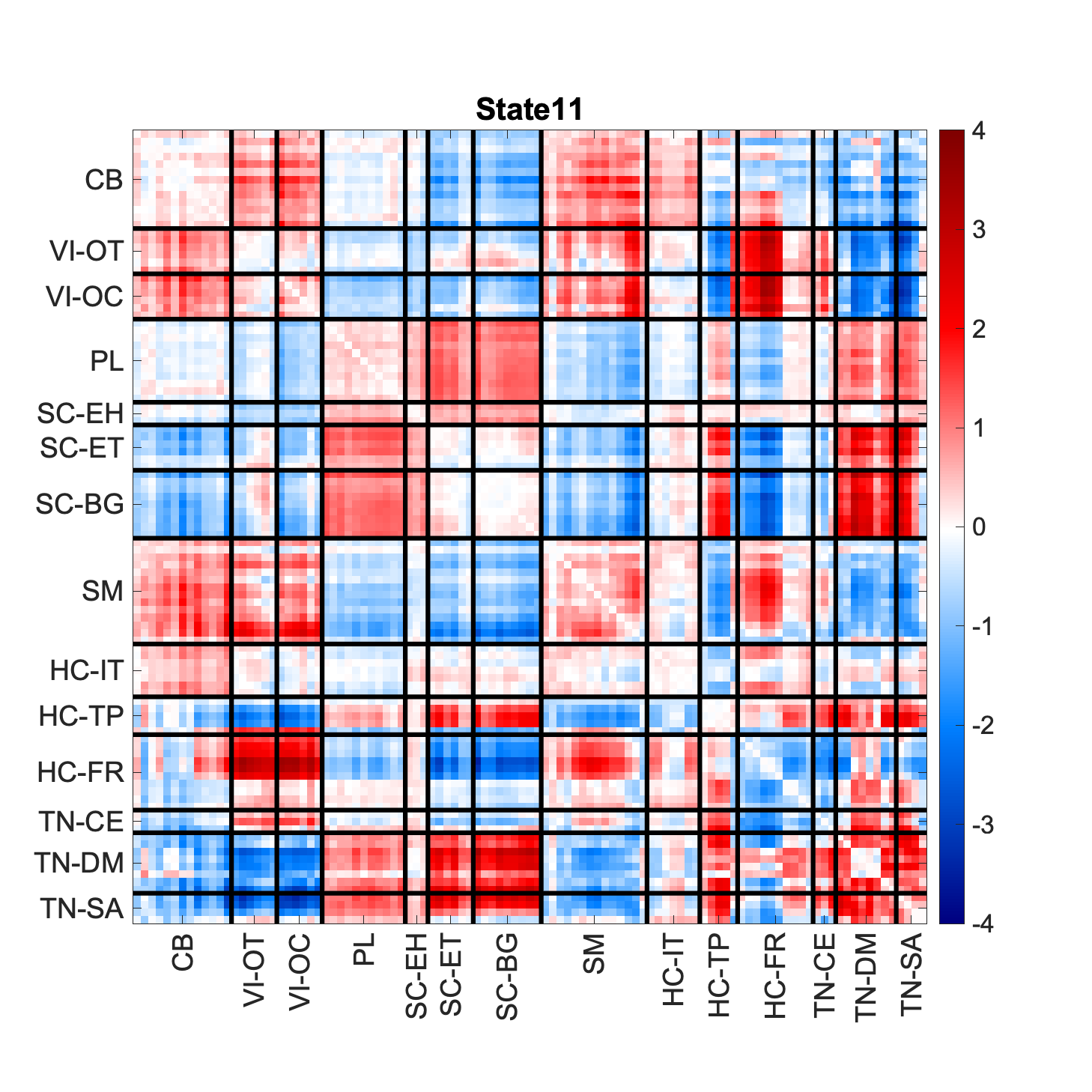

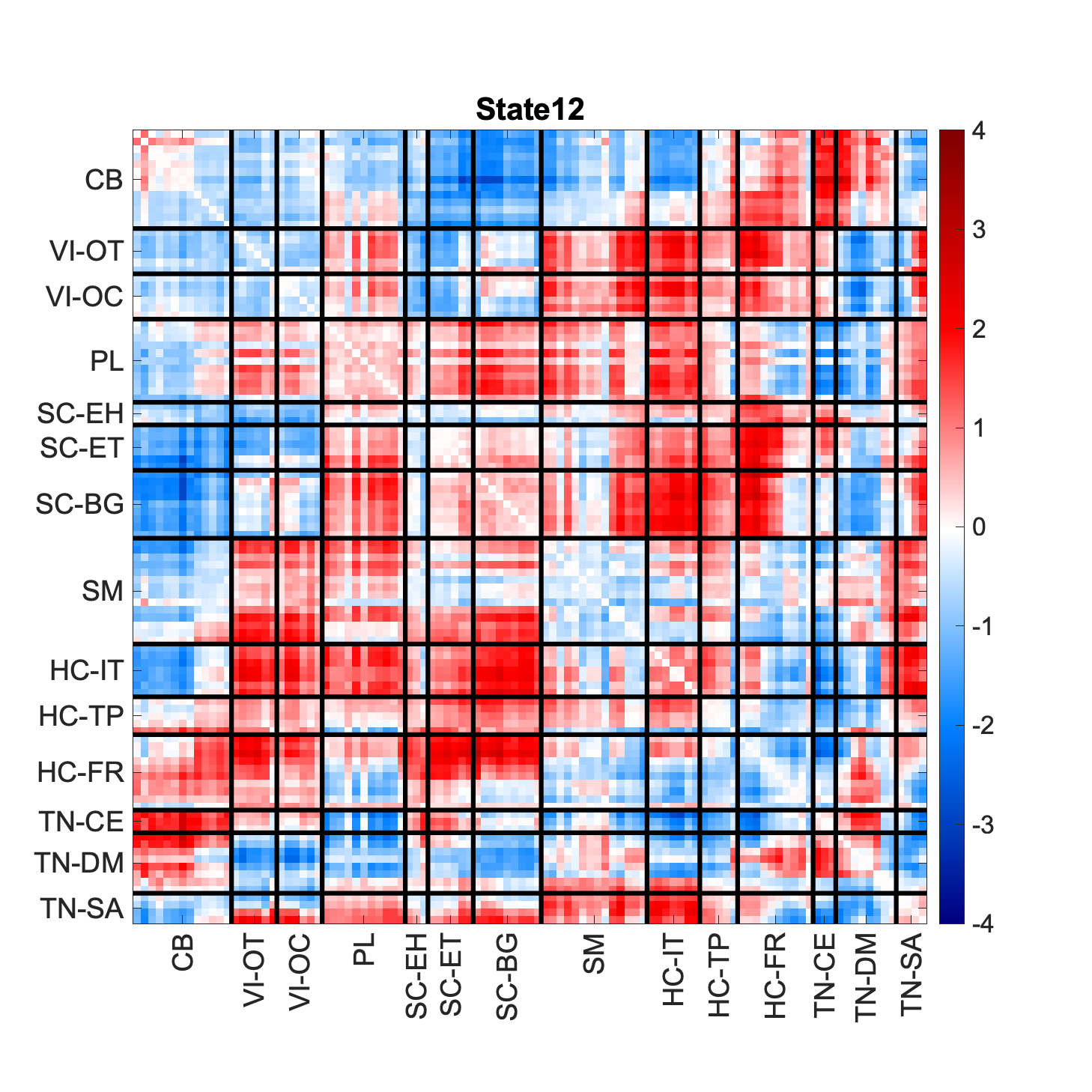

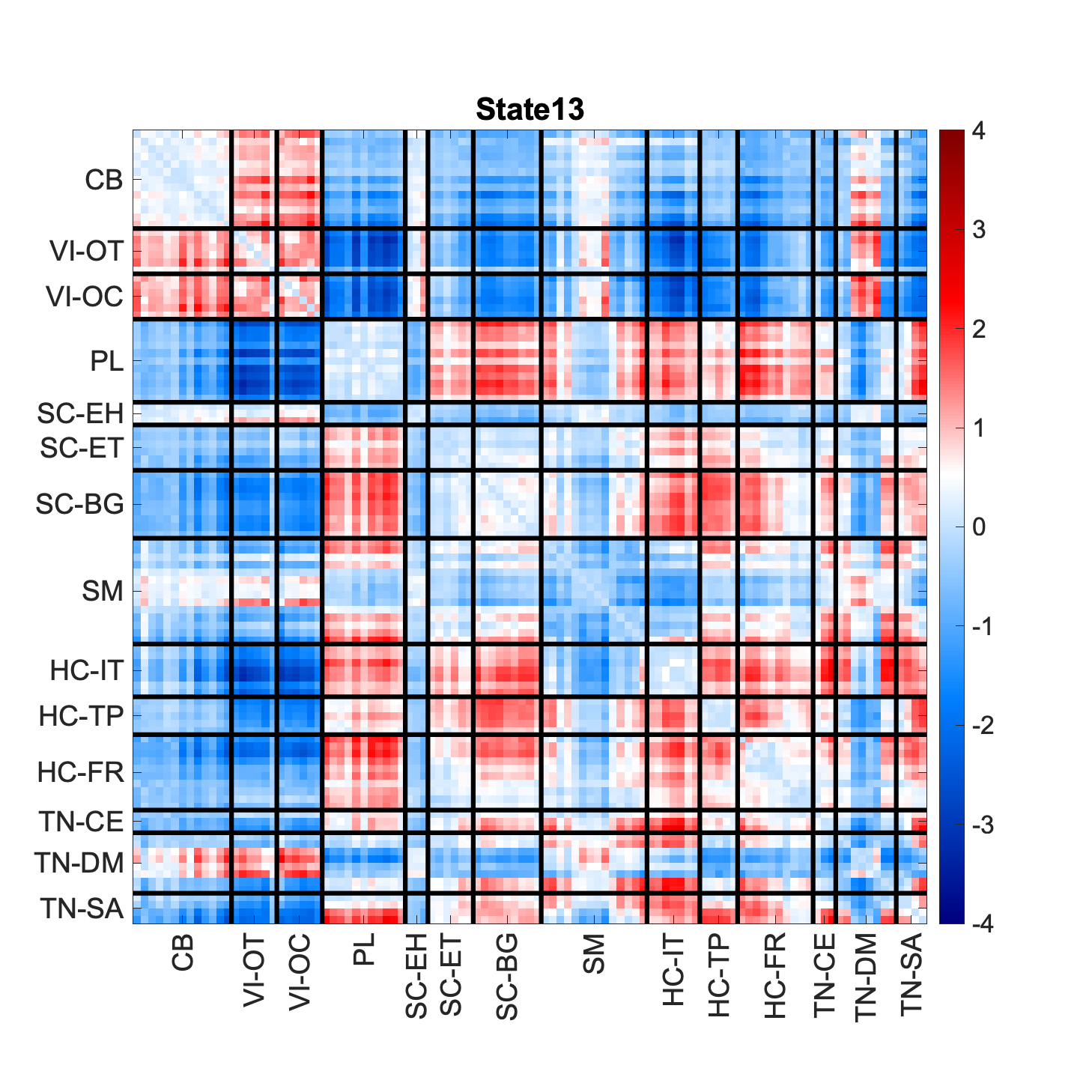

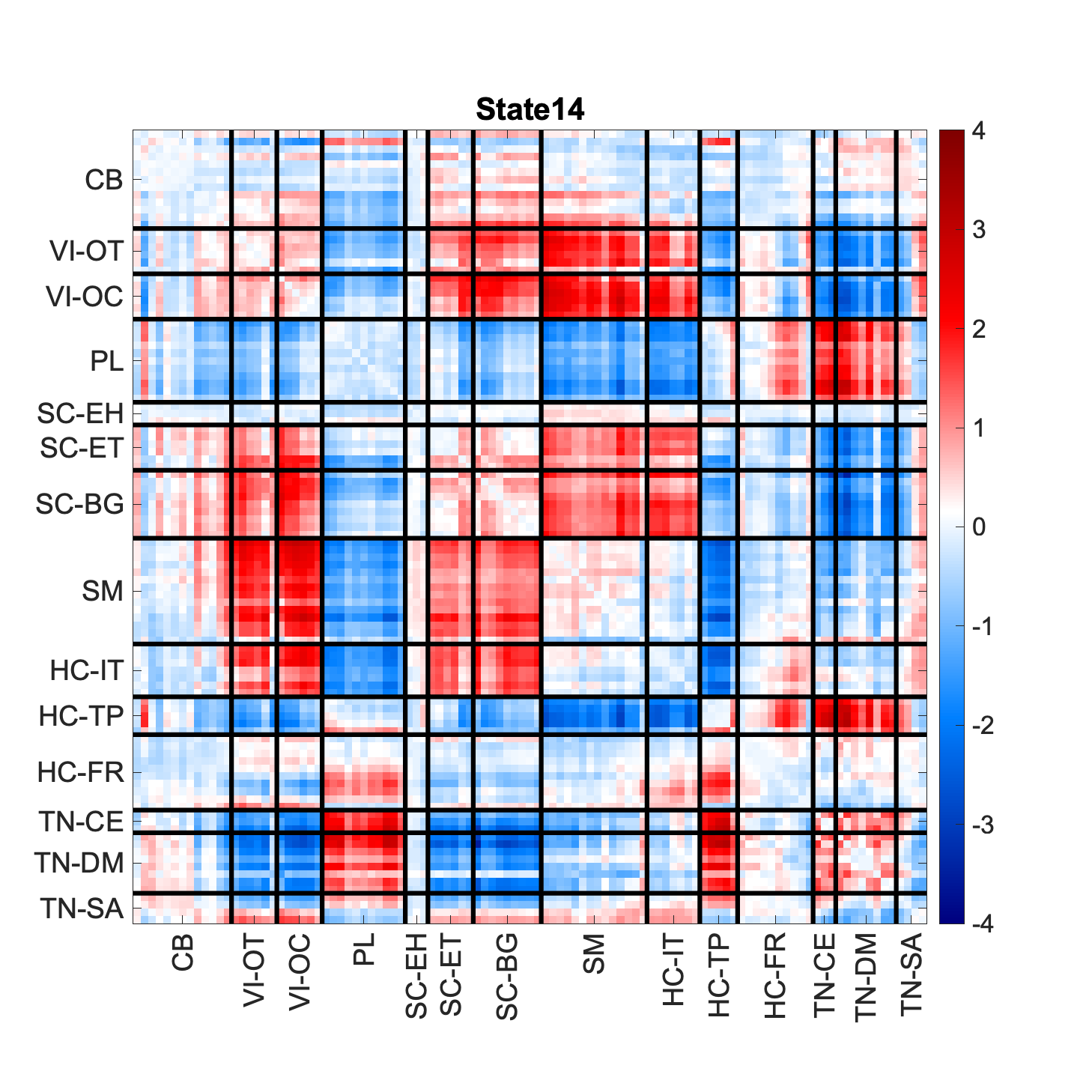

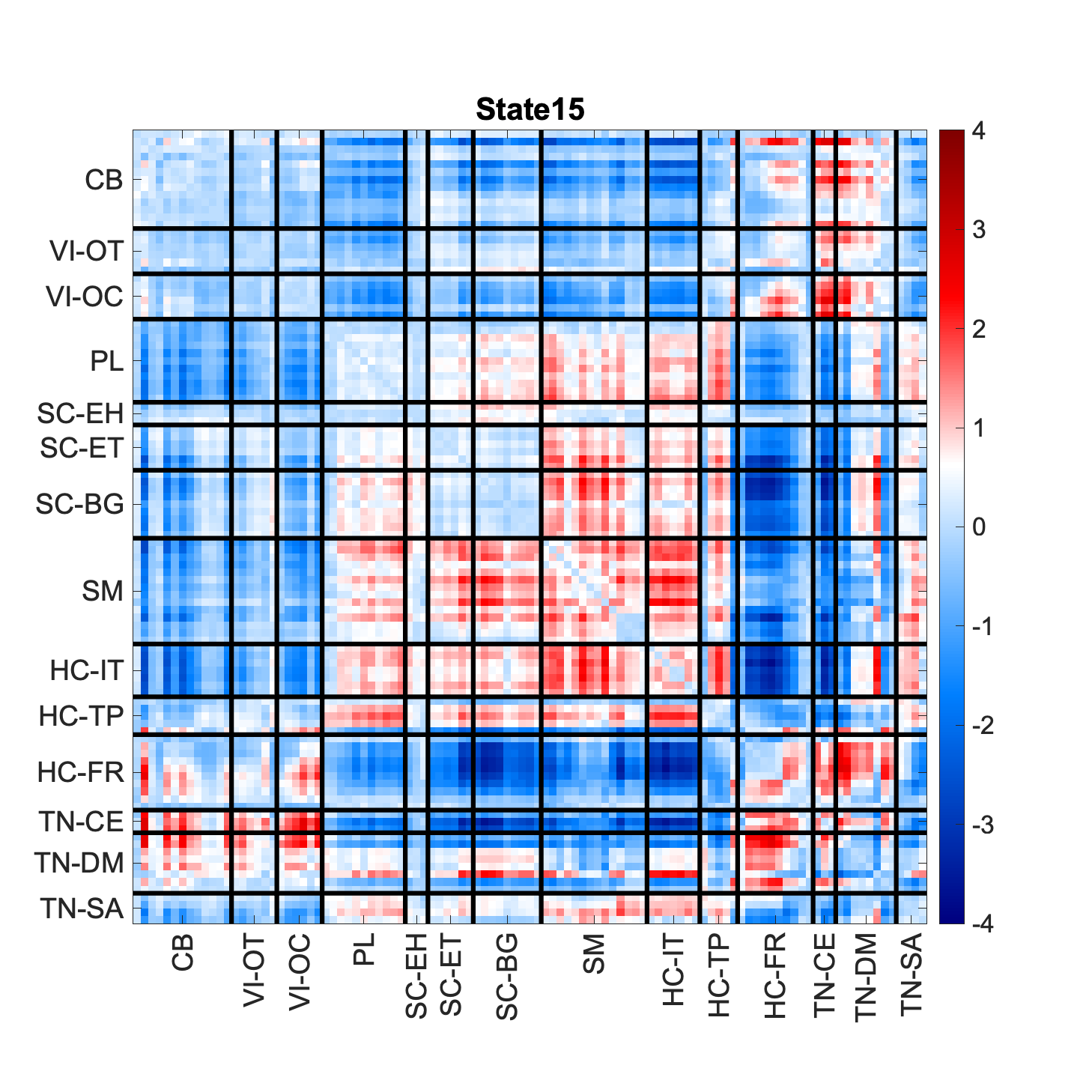

**Supplemental Figure 1:** The 15 population-level functional network connectivity (FNC) patterns or “states”. Regions in red and blue indicate positive correlation and anti-correlation between intrinsic connectivity networks (ICNs) for a specific brain state.
